# HCN channels modulate the medium afterhyperpolarization and adjust the firing gain of fast alpha motoneurons in mice

**DOI:** 10.64898/2026.05.19.726318

**Authors:** Simon A. Sharples, Gareth B. Miles

**Author notes:** Corresponding Authors **Correspondence:**, Simon A. Sharples, Division of Neurosurgery, Cincinnati Children’s Hospital, Cincinnati, Ohio, United States, 45229, Prof. Gareth B. Miles, School of Psychology and Neuroscience, University of St Andrews, Fife, United Kingdom, KY16 9JP.

## Abstract

Motoneuron subtypes exhibit distinct firing properties that are critical for the graded control of muscle force. A key determinant of these differences is the medium afterhyperpolarization (mAHP), which shapes discharge rate and firing gain. While subtype-specific variation in mAHP properties has traditionally been attributed to differences in small-conductance calcium-activated potassium (SK) channel expression, emerging evidence suggests that additional conductances may contribute. Here, we investigated the role of hyperpolarization-activated cyclic nucleotide-gated (HCN) channels in regulating the mAHP and excitability of mouse spinal motoneurons during postnatal development. Using whole-cell patch-clamp recordings, we show that, by the onset of the third postnatal week, an h current (Ih) is active at resting potential in fast motoneurons and is correlated with the amplitude of the mAHP. Pharmacological blockade of HCN channels with ZD7288 increased mAHP amplitude in fast but not slow motoneurons, without affecting mAHP duration, indicating a subtype-specific contribution to mAHP amplitude. In line with the mAHP regulating firing gain, ZD7288 also reduced firing gain in fast but not slow motoneurons. These findings support a contribution of HCN channel activity to the regulation of mAHP amplitude and firing gain in fast motoneurons, highlighting a potential interaction between Ih and SK channel-dependent mechanisms in shaping motoneuron excitability.

**Key Points:**

- The amplitude of the medium afterhyperpolarization (mAHP) is negatively correlated with h-current (Ih) amplitude measured near resting potential in mouse lumbar motoneurons.
- Pharmacological blockade of HCN channels selectively increases mAHP amplitude in fast, delayed firing alpha motoneurons, with no effect observed in slow, immediate firing alpha motoneurons.
- Inhibition of HCN channels reduces firing gain in fast motoneurons, while slow motoneurons remain unaffected.
- HCN channels regulate firing gain in fast motoneurons, at least in part, through modulation of mAHP amplitude.

## Introduction

Motor output is regulated by both the orderly recruitment and the firing rate of motoneurons. These parameters collectively define motor unit gain and can be tuned to adjust the precision or vigor of movement. Recruitment is determined by a combination of passive and active intrinsic properties, which differ systematically across motoneuron subtypes (Burke et al. 1973; Cope and Clark 1991; Sharples and Miles 2021). In contrast, firing gain is strongly influenced by voltage-gated ion channels that shape action potential discharge and afterpotentials, which also differ across motoneuron subtypes (Baldissera and Gustafsson 1971; Barrett et al. 1980; Heckman et al. 2008; Kernell 1999).

Among these properties, the medium afterhyperpolarization (mAHP) is a key determinant of firing behavior, particularly at the lower end of the frequency–current relationship, where most force is generated by motor units (Manuel et al. 2005). The mAHP constrains repetitive firing and is a defining feature that distinguishes fast and slow motoneurons (Martínez-Silva et al. 2026; Zengel et al. 1985). Differences in mAHP amplitude and duration across motoneurons subtypes have traditionally been attributed to subtype-specific expression of small-conductance calcium-activated potassium (SK) channel subunits (Deardorff et al. 2013, 2021; Dukkipati et al. 2018). However, emerging evidence challenges this view (Kissane et al. 2022), indicating that additional conductances also contribute to shaping the mAHP of motoneuron subtypes and that SK channels do not act in isolation.

Hyperpolarization-activated cyclic nucleotide-gated (HCN) channels represent a strong candidate for such modulation. HCN channels produce a hyperpolarization activated inward current (Ih) that has been classically linked to pacemaker properties and rebound bursting in neurons (Angstadt and Calabrese 1989; Chalif et al. 2022; Cymbalyuk et al. 2002; Lüthi and McCormick 1998; Siu et al. 2006; Thoby-Brisson et al. 2000; Zhu et al. 2016). However, in some neuronal subtypes, HCN channels are active at resting membrane potential, and can influence neuronal excitability through both depolarizing and shunting effects (Bayliss et al. 1994; Picton et al. 2018). For example, in hippocampal CA1 neurons, Ih has been shown to interact with afterhyperpolarization mechanisms, altering their amplitude and functional impact on firing gain (Dwivedi and Bhalla 2021; Gu et al. 2005). Consistent with this notion, classic studies in cat motoneurons reported a correlation between the magnitude of the sag potential, an indicator of Ih, and the mAHP (Gustafsson and Pinter 1984). However, a direct role for HCN channels in regulating the mAHP and firing gain in spinal motoneurons has not been established.

We have previously demonstrated that a resting h-current emerges selectively in fast motoneurons at the onset of the third postnatal week in mice. This observation raises the possibility that HCN channel activity may contribute to subtype-specific differences in mAHP properties and firing gain. Here, we test this hypothesis using a combination of reanalysis of published datasets and new, targeted pharmacological experiments. Our findings support a novel mechanism in which HCN channels shape the amplitude of the mAHP and regulate firing gain in fast motoneurons.

## Results

We examined intrinsic properties of motoneuron subtypes distinguished by their firing responses to 5-s depolarizing current steps. Fast α-motoneurons were identified by a delayed onset to repetitive firing and an accelerating discharge, whereas slow motoneurons displayed immediate firing with stable or adapting discharge rates (Leroy et al. 2014). Consistent with prior work (Leroy et al. 2014; Martínez-Silva et al. 2026; Sharples and Miles 2021; Zengel et al. 1985), delayed firing motoneurons (n = 11) exhibited a medium afterhyperpolarization (mAHP) that was smaller in amplitude (Fig. 1C, D; t(17) = 4.6, p = 0.0003) and shorter in duration (Fig. 1E; t(17) = 3.8, p = 0.002) compared with immediate firing motoneurons (n = 8).

**Figure 1:**
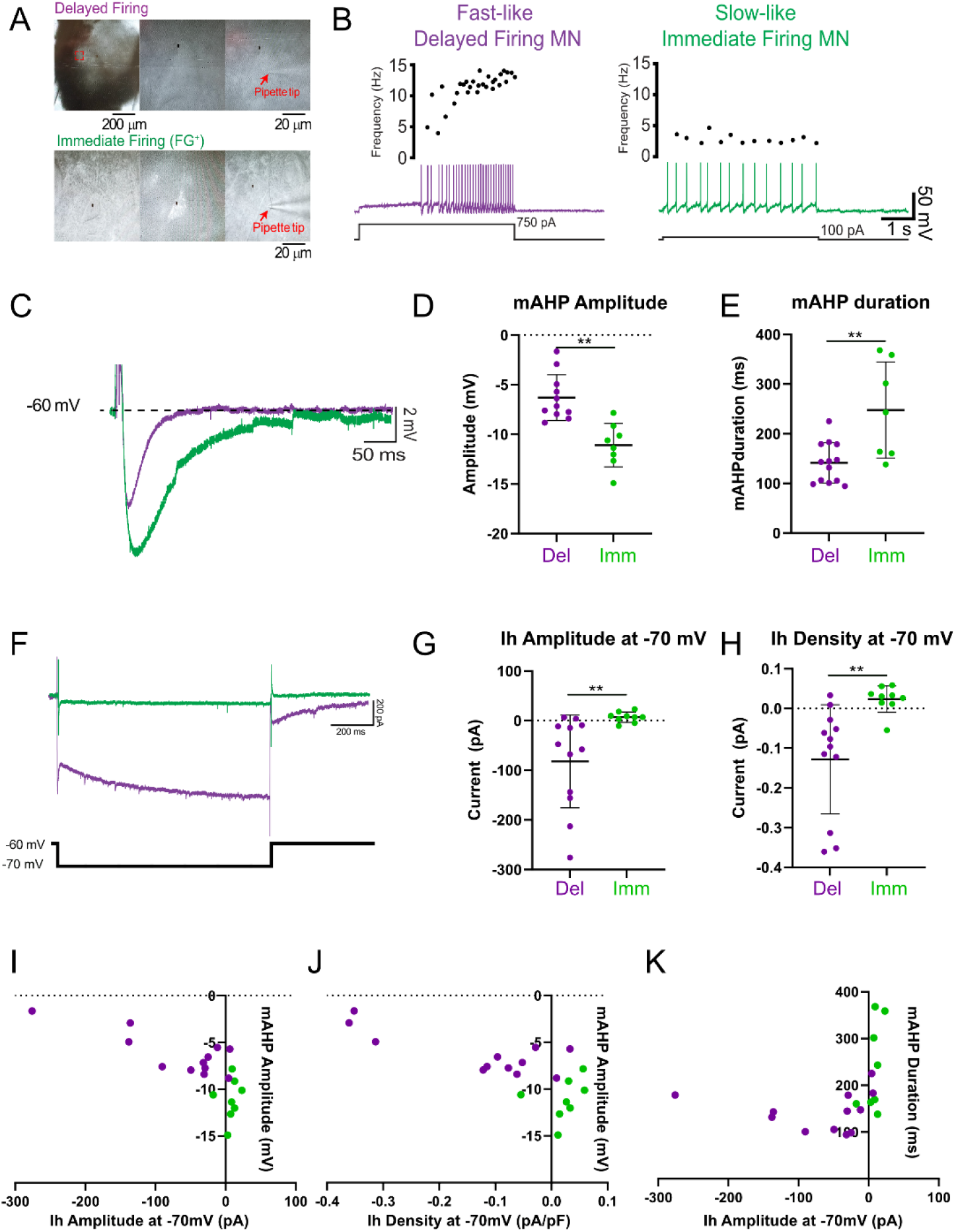
A smaller medium afterhyperpolarization (mAHP) in delayed firing motoneurons is correlated with a resting h current. (A) Motoneurons visualized under differential interference contrast were identified based on delayed (purple) and immediate (green) firing in response to long (5 second) depolarizing current steps applied near rheobase current (B). A-B, adapted from (Sharples and Miles 2021). C) The mAHP of a delayed (purple) and immediate (green) firing motoneuron following a 10 ms depolarizing current step applied at an intensity of 2 times rheobase current (not shown). D) Delayed firing motoneurons (n = 11) have a larger amplitude (D; p = 0.0003) and longer duration (E; p = 0.002) mAHP compared to immediate firing motoneurons (n = 7). F) The h current (Ih) was measured near resting potential in delayed and immediate firing motoneurons in voltage clamp in response to a 10 mV hyperpolarizing voltage step from a holding potential of −60 mV. Delayed firing motoneurons present with an h-current near resting potential (−70 mV) that is not found in immediate firing motoneurons (G; p = 0.01). H) The density of the h-current at −70 mV is larger in delayed compared to immediate firing motoneurons (p = 0.005). Scatter plot between the amplitude of the mAHP and the h current amplitude (I; r = −0.7, p = 0.003) and density (J; r = −0.7, p = 0.0002). The duration of the mAHP was not correlated with the amplitude of the h current (K; r = 0.3, p = 0.2). Data are presented as individual data points and were analyzed using an unpaired t-test. Asterisks are displayed and p values reported where significant differences were detected.

Given that the h current (Ih) is active near resting membrane potentials in delayed firing motoneurons (Sharples and Miles 2021), we tested the hypothesis that Ih helps shape the mAHP by opposing the hyperpolarizing actions of the mAHP. Delayed firing motoneurons exhibited larger h current amplitudes at −70 mV (Fig. 1F, G; t(17) = 2.8, p = 0.01) and greater Ih density (Fig. 1H; t(17) = 3.2, p = 0.005) than immediate firing motoneurons, consistent with previous work (Sharples and Miles 2021; Sharples and Miles, 2026). Across all cells, h current amplitude measured near resting potential at −70 mV was inversely correlated with mAHP amplitude (Fig. 1I; r = −0.73, p = 0.0003), supporting the idea that greater Ih is associated with reduced mAHP magnitude. This relationship persisted when normalized for cell size using Ih density (Fig. 1J; r = −0.73, p = 0.0002). In contrast, Ih amplitude was not correlated with mAHP duration (Fig. 1K; r = 0.3, p = 0.14), suggesting a selective influence on mAHP amplitude rather than kinetics.

To directly test this hypothesis, we pharmacologically blocked HCN channels using ZD7288 (10 uM, n = 11 delayed, n = 7 immediate). This manipulation abolished Ih (Fig. 2A, B (F(1,16) = 8.5, p = 0.009). Consistent with the correlative findings, HCN channel blockade increased the mAHP amplitude in delayed firing motoneurons by 34±44%, with no effect in immediate firing motoneurons (−3±34%; Figure 2C, D; F(1,16) = 5.3, p = 0.03). The mAHP duration was unchanged by ZD7288 in either subtype (del: 25±40%, imm: −1±29%; Figure 2E; F(1,16) = 0.9, p = 0.4).

**Figure 2:**
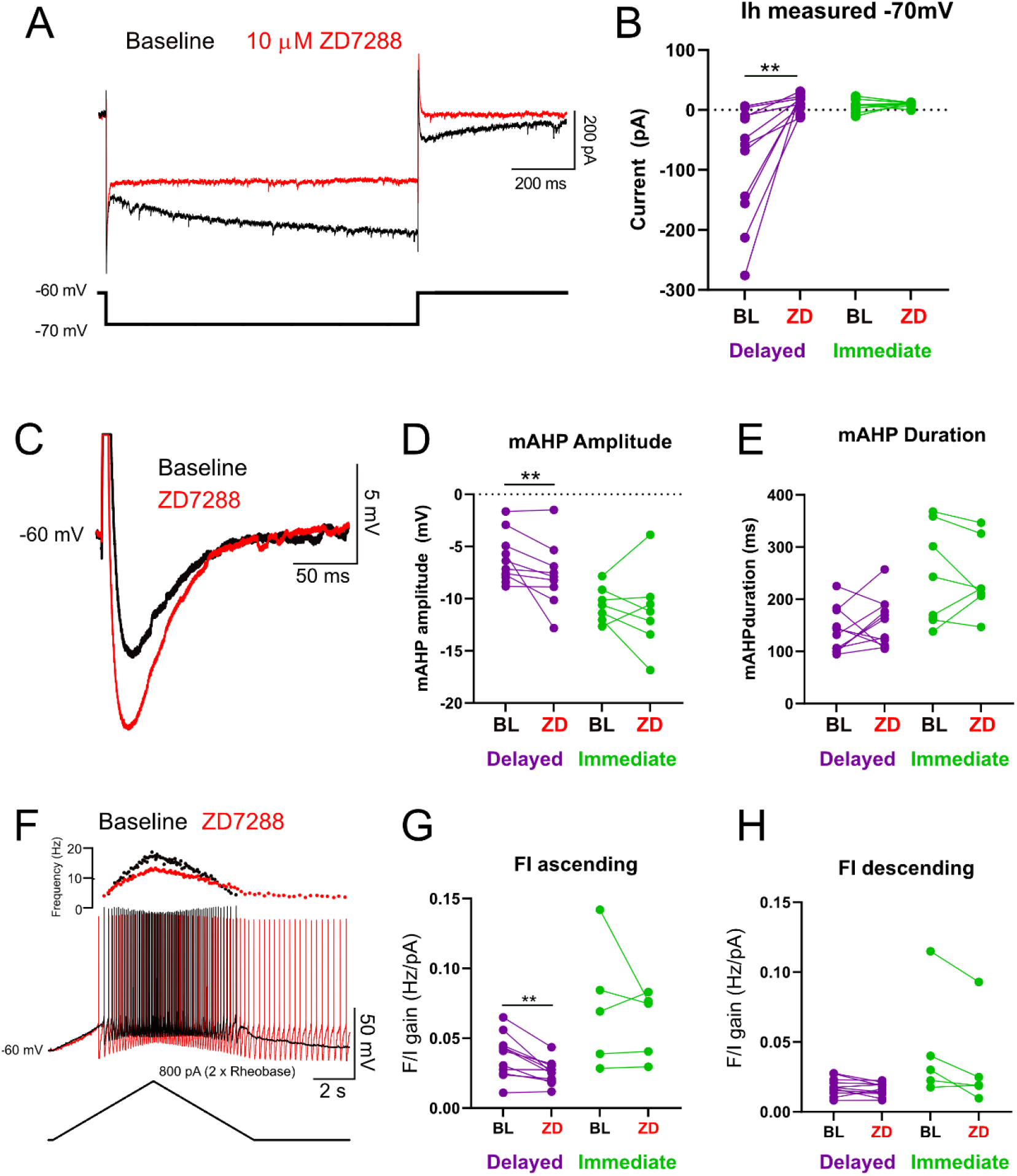
Blocking HCN channels increases the amplitude of the mAHP and reduced firing gain in delayed firing motoneurons. (A) Representative voltage clamp traces illustrating the h-current (Ih) measured near resting potential in response to a 10-mV hyperpolarizing voltage step applied from −60 mV before (black) and after application of the HCN channel blocker ZD7288 (10 µM) in 11 delayed firing motoneurons and 7 immediate firing motoneurons obtained from mice during the third postnatal week (P14-20). (B) ZD7288 eliminates the h current measured at −70 mV in delayed firing motoneurons (p = 0.008). (C) Representative traces of the mAHP of a delayed firing motoneuron before (black) and after application of the HCN channel blocker ZD7288 (Red; 10 uM). ZD7288 increased the amplitude of delayed (p = 0.01) but not immediate firing (p = 0.6) motoneurons (D) but did not alter mAHP duration in either motoneuron subtype (E). (F) Repetitive firing in response to a depolarizing triangular current ramp applied to an intensity of 2 times rheobase in a delayed firing motoneuron before (black) and after application of ZD7288. Frequency-current (F/I) gain, measured as the slope, on the ascending (p = 0.03) (G) and descending limb (H) of triangular current ramp before and after ZD7288 in delayed and immediate firing motoneurons. Data are presented as individual data points and were analyzed using a two-factor repeated measures ANOVA. Asterisks are displayed and p values reported when significant differences were detected using a Holm-Sidak post hoc test.

Because the mAHP is a key determinant of spike-frequency adaptation and firing gain (Powers et al. 1999), we next assessed firing responses using triangular depolarizing current ramps (2× rheobase) (Figure 2F). This approach enabled comparison of firing behavior during the ascending and descending phases of the ramp, where Ih activation is expected to differ. Given that increasing mAHP amplitude is predicted to reduce firing gain, we hypothesized that HCN channel blockade would decrease motoneuron firing during the ramps. Consistent with previous reports (Sharples et al., 2026), ZD7288 reduced the current required for recruitment (BL: 823±816 pA ZD: 525±474 pA, p = 0.004) and derecruitment (BL: 590±771 pA ZD: 245±400 pA, p = 0.03), but did not alter the maximal firing rate attained during the ramp (BL: 19.2±8.4 ZD:, 21.2±6.3 Hz; p = 0.3). Interesting, and in line with our hypothesis, blocking HCN channels reduced firing gain in delayed firing motoneurons, with minimal effect in immediate firing motoneurons. This effect was evident on the ascending limb of the ramp (Figure 2G; F(1,14) = 5.4, p = 0.03), whereas firing on the descending limb was largely unchanged (Figure 2H; F(1,14) = 2.1, p = 0.2). The lack of effect during the descending phase indicated the differential effect of HCN channel blockade on firing behaviour during recruitment versus derecruitment.

## Discussion

The present study suggests that hyperpolarization-activated cyclic nucleotide-gated (HCN) channels shape the medium afterhyperpolarization (mAHP) and differentially regulate firing gain in spinal motoneuron subtypes. In line with previous work (Gustafsson and Pinter 1984; Leroy et al. 2014; Martínez-Silva et al. 2026; Rotterman et al. 2021; Sharples and Miles 2021; Zengel et al. 1985), we show that fast α-motoneurons exhibit a smaller mAHP than slow motoneurons but also uniquely express a resting hyperpolarization-activated inward current (Ih). Pharmacological blockade of HCN channels with ZD7288 abolishes this resting Ih, increases mAHP amplitude, and reduces firing gain selectively in fast motoneurons. These findings are consistent with a contribution of HCN channel activity to the regulation of mAHP amplitude, which in turn is known to influence motoneuron firing gain. Here, we discuss these findings in the context of previous work on motoneuron intrinsic properties, neuromodulatory control, and dysfunction in amyotrophic lateral sclerosis (ALS).

The mAHP is a defining electrophysiological feature that differentiates motoneuron subtypes and plays a central role in determining discharge rate and firing gain (Kernell and Zwaagstra 1981). Here, we present evidence that is consistent with HCN channels contributing to the amplitude of the mAHP in a motoneurons subtype-specific manner, providing a mechanism that complements existing SK channel-based frameworks. Traditionally, differences in mAHP properties have been attributed to variations in small-conductance calcium-activated potassium (SK) channel expression and kinetics (Abdul Halim et al. 2026; Baldissera and Gustafsson 1974; Berger et al. 1995; Miles et al. 2005). Consistent with this view, previous work has shown that a subset of smaller α-motoneurons with slower conduction velocities, corresponding to putative slow motoneurons, exhibit larger and longer-lasting mAHPs and express both SK2 and SK3 subunits. In contrast, the larger motoneurons, with faster conduction velocities, corresponding to putative fast motoneurons, had smaller and shorter-lasting mAHPs but only expressed SK3 subunits (Deardorff et al. 2013). These SK channel subunits are enriched at C-bouton synapses, forming a key substrate for cholinergic modulation of motoneuron excitability. However, more recent work has demonstrated that SK3 expression is not restricted to motoneurons innervating slow-twitch muscle fibers (Kissane et al. 2022), indicating that SK subunit distribution alone cannot fully account for subtype-specific differences in mAHP properties. In addition, these studies primarily examined clustering of SK2 and SK3 subunits at C-bouton synapses, whereas SK channels located outside of these synaptic domains are also likely to contribute to shaping the mAHP (Mousa and Elbasiouny 2020; Venugopal et al. 2012). In this context, our findings are consistent with HCN channels contributing to the regulation of the mAHP. Specifically, Ih was strongly correlated with mAHP amplitude, and pharmacological blockade of HCN channels increased mAHP amplitude in fast motoneurons. In contrast, mAHP duration was not correlated with Ih measured near resting potential, and HCN channel blockade did not alter mAHP duration. This dissociation suggests that HCN channels may primarily regulate mAHP amplitude, whereas other conductances, including SK channels, may play a more prominent role in determining its duration. These results build on our previous observations that a resting Ih emerges selectively in fast motoneurons at the onset of the third postnatal week (Sharples and Miles 2021). This developmental shift in the activation range of Ih coincides with the maturation of subtype-specific intrinsic properties, supporting a functional role in establishing excitability differences between motoneuron subtypes. Here, we extend these findings by showing that, in addition to influencing recruitment, Ih may also actively modulate mAHP amplitude and, consequently, firing gain in fast motoneurons. The observed correlation between Ih magnitude and mAHP amplitude further supports this relationship and is consistent with earlier work in cat motoneurons (Gustafsson and Pinter 1984), as well as studies in other neuronal types, including hippocampal CA1 and cortical neurons (Dwivedi and Bhalla 2021; Gu et al. 2005). We therefore propose that depolarizing inward current produced by HCN channels during the mAHP reduces its effective amplitude, thereby increasing firing gain in fast motoneurons.

The functional implications of this mechanism are particularly relevant in the context of neuromodulation. Motoneuron firing gain is dynamically regulated by a range of neuromodulatory inputs, many of which converge on the mAHP (Miles and Sillar 2011; Perrier et al. 2013; Rekling et al. 2000; Sharples et al. 2014) (Miles and Sillar, 2011; Rekling, et al., 2000; Perrier, et al., Shaples, et al., 2014). For example, serotonin (Perrier and Cotel 2015), acetylcholine (Miles et al. 2007; Revill et al. 2019), and dopamine (Han et al. 2007; Sharples et al. 2020) all increase motoneuron firing gain while reducing mAHP amplitude. More recently, muscarinic receptor activation has been shown to selectively reduce mAHP amplitude and enhance firing gain in fast motoneurons (Eleftheriadis et al. 2023). These effects have traditionally been attributed to modulation of SK channel function. However, in light of the present findings, an additional mechanism involving HCN channels should be considered. HCN channels are well-established targets of neuromodulators, and their activation properties are strongly regulated by intracellular signaling pathways, including cAMP (Wainger et al. 2001). In other neuronal systems, neuromodulators can shift the voltage dependence of HCN channels, thereby altering the contribution of Ih at subthreshold membrane potentials (McCormick and Pape 1990; Peck et al. 2006; Robinson and Siegelbaum 2003). Similarly, serotonin depolarizes the activation range of Ih in neonatal motoneurons, increasing its contribution near resting membrane potentials (Bayliss et al. 1997; Kjaerulff and Kiehn 2001). In this context, modulation of HCN channel activity is well positioned to influence mAHP amplitude through a shunting or depolarizing mechanism. Our findings therefore suggest that neuromodulators may regulate firing gain not only through SK channel-dependent mechanisms, but also by tuning Ih, particularly in fast motoneurons where a resting Ih is present. This provides a potential framework for subtype-specific control of motoneuron excitability via convergent modulation of multiple conductances that shape the mAHP.

These findings may also have important implications for motoneuron disease, where dysregulation of ion channel function contributes to aberrant excitability and motoneuron degeneration (Bączyk et al. 2022; LoRusso et al. 2019; Odierna et al. 2024; Stringer and Weiss 2023). In amyotrophic lateral sclerosis (ALS), alterations in motoneuron intrinsic properties evolve over disease progression and differ across motoneuron subtypes and mouse models (Delestrée et al. 2014; Leroy et al. 2014; Martínez-Silva et al. 2018; Pambo-Pambo et al. 2009; Quinlan et al. 2011). Notably, consistent with the effects of HCN channel blockade observed here, motoneurons at symptomatic stages in SOD1-G93A mice exhibit an increased mAHP amplitude accompanied by a reduced sag ratio, suggesting a loss of Ih contribution (Jensen et al. 2020). These changes are not evident at presymptomatic stages (Delestrée et al. 2014; Leroy et al. 2014; Quinlan et al. 2011), indicating that disruption of HCN channel function may emerge with disease progression. In contrast, inducible TDP-43 models display reduced mAHP amplitude and increased frequency–current (F– I) gain, without a corresponding change in sag amplitude (Djukic et al. 2025), suggesting that enhanced excitability in this context arises independently of alterations in Ih. Together, these findings highlight substantial variability across ALS models, with divergent changes in mAHP properties, Ih-related conductances, and firing gain. While some studies report increased excitability, others demonstrate that fast motoneurons become hypoexcitable at later stages of disease (Devlin et al. 2015; Martínez-Silva et al. 2018), underscoring the dynamic and stage-dependent nature of these changes. This variability is consistent with the engagement of homeostatic mechanisms that act to stabilize motoneuron output in the face of ongoing degeneration (Huh et al. 2021; Nascimento et al. 2024). In this framework, compensatory adjustments in multiple conductances—including SK channels, HCN channels, and other intrinsic currents—may differentially shape mAHP amplitude and firing gain across disease models and stages. Our results therefore raise the possibility that alterations in HCN channel function, and their interaction with mAHP-generating mechanisms could contribute to the shifting motoneuron excitability observed over the course of ALS progression.

Several limitations should be considered when interpreting these findings. First, although pharmacological blockade with ZD7288 is widely used to inhibit HCN channels, off-target effects on membrane and synaptic properties have been reported (Felix et al. 2003; Wu et al. 2012), and thus we cannot fully exclude indirect contributions to the observed changes in mAHP amplitude and firing gain. However, given that ZD7288 did not affect the mAHP amplitude of slow motoneurons, despite being applied under identical experimental conditions, it is unlikely that the observed effects in fast motoneurons were driven primarily by nonspecific off-target actions. Rather, these findings support the interpretation that the changes in mAHP amplitude and firing gain predominantly reflect blockade of HCN channel-mediated conductances. Second, the relationship between Ih and mAHP amplitude is supported in part by correlative analyses, and while the pharmacological data are consistent with a functional contribution of HCN channels, they do not establish a direct causal mechanism. Finally, motoneuron subtypes were classified based on established electrophysiological criteria, which, while widely used (Bhumbra and Beato 2018; Bos et al. 2018; Leroy et al. 2014; Nascimento et al. 2024; Özyurt et al. 2022; Sharples et al. 2023, 2025), do not provide definitive identification of motor unit type (Martínez-Silva et al. 2026). Despite these limitations, the convergence of correlative and pharmacological findings, together with the subtype-specific effects observed, supports a model in which HCN channel activity contributes to the regulation of mAHP amplitude and firing gain in fast motoneurons.

In summary, our findings provide converging evidence that HCN channel activity contributes to the regulation of mAHP amplitude and firing gain in fast motoneurons. By attenuating the mAHP, a resting Ih is well positioned to represent an additional mechanism through which intrinsic conductances shape subtype-specific excitability. These results extend existing frameworks centered on SK channel-dependent processes and highlight HCN channels as a complementary component in the control of motoneuron output.

## Additional Information

### Contributions

Study Conception and Design: SAS, GBM; Data acquisition and analysis: SAS; Preparation of Figures: SAS; First draft and revision of manuscript: SAS, GBM; All authors approved the final version of the manuscript.

## Acknowledgements

The authors reserve the right to apply a Creative Commons Attribution (CC BY) license to any Author Accepted manuscript version arising from this submission.

## Data Availability Statement

The research data supporting this publication will be made freely available in an open access data repository following acceptance to a peer-reviewed journal.

## Methods

### Animals

This study included unpublished data in addition to reanalysis of experimental data summarized in (Sharples et al. 2025; Sharples and Miles 2021; Sharples and Miles, 2026), which included experiments performed on tissue obtained from 17 (male: n = 9; and female: n = 8) wild type C57Bl/6J mice at postnatal days (P) 14-20. All procedures were conducted in accordance with the UK Animals (Scientific Procedures) Act 1986. Experiments were approved by the University of St Andrews Animal Welfare Ethics Committee and were covered under a Project License (PP8253850) approved by the Home Office. All animals were provided with unrestricted access to food and water and housed in climate-controlled conditions.

### Tissue preparation

Animals were sourced from an in-house colony within the St Mary’s Animal Unit at the University of St Andrews. Animals were killed using Schedule 1 procedures defined by the Home Office by performing a cervical dislocation followed by rapid decapitation. Animals were then eviscerated and pinned ventral side up in a dissecting chamber lined with silicone elastomer (Sylguard), filled with ice-cold (1-2 degrees Celsius) potassium gluconate based dissecting/slicing aCSF (containing in mM: 130 K-gluconate, 15 KCl, 0.05 EGTA, 20 HEPES, 25 D-glucose, 3 kynurenic acid, 2 Na-pyruvate, 3 myo-inositol, 1 Na-L-ascorbate; pH 7.4, adjusted with NaOH; osmolarity approximately 345 mOsm) that was continuously bubbled with carbogen (95% oxygen, 5% carbon dioxide). Spinal cords were exposed by performing a ventral vertebrectomy, cutting the dorsal roots and gently lifting the spinal cord from the spinal column. Spinal cords were removed within 3 - 5 minutes following cervical dislocation. Spinal cords were secured directly to an agar block (3 % agar) with VetBond surgical glue (3M) and glued to the base of the slicing chamber with cyanoacrylate adhesive. The tissue was immersed in ice-cold dissecting/slicing aCSF and bubbled with carbogen. Blocks of frozen slicing solution were also placed in the slicing chamber to keep the solution around 1-2 degrees Celsius. On average, the first slice was obtained within 10 minutes of decapitation which increased the likelihood of obtaining viable motoneurons in slices. 300 µm transverse slices were cut at a speed of 10 um/s on the vibratome (Leica VT1200) to minimize tissue compression during slicing. 3-4 slices were obtained from each animal. Slices were transferred to a recovery chamber filled with pre-warmed (35 degrees Celsius) recovery aCSF (containing in mM: 119 NaCl, 1.9 KCl, 1.2 NaH2PO4, 10 MgSO4, 1 CaCl2, 26 NaHCO3, 20 glucose, 1.5 kynurenic acid, 3% dextran), continuously bubbled with carbogen, for thirty minutes after completion of the last slice, which took 10 - 15 minutes on average. Following recovery, slices were transferred to a chamber filled with warm (35 degrees Celsius) recording aCSF (containing in mM: 127 NaCl, 3 KCl, 1.25 NaH2PO4, 1 MgCl2, 2 CaCl2, 26 NaHCO3, 10 glucose), bubbled with carbogen, and allowed to equilibrate at room temperature (maintained at 23-25 degrees Celsius) for at least one hour before experiments were initiated.

### Whole cell patch clamp electrophysiology

This study includes data from whole-cell patch clamp recordings obtained from a total of 22 (13 fast, 9 slow) lumbar motoneurons, from which baseline mAHP measurements included in (Sharples et al. 2025; Sharples and Miles 2021), h current measurements before and after ZD7288 were included in (Sharples and Miles 2021), and firing gain measured frequency-current plots during triangular depolarizing current ramps in (Sharples and Miles 2026). In these experiments, spinal cord slices were stabilized in a recording chamber with fine fibers secured to a platinum harp and visualized with a 40x objective with infrared illumination and differential interference contrast (DIC) microscopy. A large proportion of the motoneurons studied were identified based on location in the ventrolateral region with somata greater than 20 µm. Recordings were obtained from a subset of motoneurons that had been retrogradely labelled with Fluorogold (Fluorochrome, Denver, CO). Fluorogold was dissolved in sterile saline solution and 0.04 mg/g injected intraperitoneally 24-48 hours prior to experiments (Miles et al. 2005). In addition to recording from larger FG-positive cells, this approach allowed us to more confidently target smaller motoneurons. Motoneurons were visualized and whole cell recordings obtained under DIC illumination with pipettes (L: 100 mm, OD: 1.5 mm, ID: 0.84 mm; World Precision Instruments) pulled on a Flaming Brown micropipette puller (Sutter instruments P97) to a resistance of 2.5-3.5 MΩ. Pipettes were back-filled with intracellular solution (containing in mM: 140 KMeSO4, 10 NaCl, 1 CaCl2, 10 HEPES, 1 EGTA, 3 Mg-ATP and 0.4 GTP-Na2; pH 7.2-7.3, adjusted with KOH).

Signals were amplified and filtered (6 kHz low pass Bessel filter) with a Multiclamp 700 B amplifier, acquired at 20 kHz using a Digidata 1440A digitizer with pClamp Version 10.7 software (Molecular Devices) and stored on a computer for offline analysis.

### Data acquisition and analysis

All parameters were studied as in (Sharples and Miles 2021). All motoneuron intrinsic properties were studied by applying a bias current to maintain the membrane potential at −60 mV. Values reported are not liquid junction potential corrected to facilitate comparisons with previously published data (Durand et al. 2015; Miles et al. 2007; Nascimento et al. 2020; Özyurt et al. 2022; Quinlan et al. 2011; Sharples and Miles 2021; Smith and Brownstone 2020). Cells were excluded from analysis if access resistance was greater than 20 MΩ or changed by more than 5 MΩ over the duration of the recording, or if spike amplitude measured from threshold was less than 60 mV. Motoneuron subtypes were identified using a protocol established by (Leroy et al. 2014), which differentiates motoneuron type based on the latency to the first spike when injecting a 5 second square depolarizing current near the threshold for repetitive firing. Using this approach, we were able to identify 2 main firing profiles - a delayed repetitive firing profile with accelerating spike frequency, characteristic of fast-type motoneurons, and an immediate firing profile with little change in spike frequency, characteristic of slow-type motoneurons (Figure 1). The medium afterhyperpolarization (mAHP) was elicited and measured following the generation of a single action potential using a 10 ms square, depolarizing current pulse applied at an intensity 2 times rheobase current. A total of 10 trials were elicited and averaged for each cell at baseline and following drug applications. The h current (Ih) was measured in voltage clamp during 1 second hyperpolarizing voltage steps from - 60 mV to - 110 mV, in 10 mV increments. Ih was measured as the difference between the initial and steady state current. Current density was calculated by normalizing current amplitude to each cells respective whole cell capacitance. Firing gain was assessed in current clamp using triangular, depolarizing current ramps with 5 second rise and fall times (Bennett et al. 2001; Durand et al. 2015; Li and Bennett 2003; Steele et al. 2020). Triangular depolarizing current ramps were set to a peak current of 2 times repetitive firing threshold current (determined with a 100pA/s depolarizing current ramp initiated from −60 mV). Firing gain was determined by measuring the slope of the frequency-current relationship during ascending and descending aspects of the ramp.

### Research design and statistical analysis

Two factor repeated measures analysis of variance (ANOVA) were performed when comparing the effect of blocking HCN channels on the mAHP or h current. Holm-Sidak post hoc analysis was performed when significant main effects of interactions were detected. Paired or unpaired t-tests were performed when comparing two conditions. Appropriate and equivalent nonparametric tests (Mann-Whitney or Kruskal-Wallis) were conducted when data failed tests of normality or equal variance with Shapiro Wilk and Brown-Forsythe tests, respectively. Individual data points for all cells are presented in figures with mean ± SD. Statistical analyses were performed using Graph Pad Version 9.0 (Prism, San Diego, CA, USA).

